# Steep logarithmic increase of genetic variation in natural *Arabidopsis thaliana* accessions across geographic scales

**DOI:** 10.1101/2024.04.26.591275

**Authors:** Vera Hesen, Yvet Boele, Rens Holmer, René Boesten, Raúl Wijfjes, Mark G. M. Aarts, Wim H. van der Putten, Ben Scheres, Viola Willemsen

## Abstract

*Arabidopsis thaliana’s* large native range across Eurasia and display of considerable genetic variation is key to its increasing use in eco-evolutionary studies. The structure and amount of this genetic variation has been studied on various geographic scales. On a continental scale, the genetic variation was postulated to follow an ‘isolation by distance’ model, implying less genetic variation at smaller geographic distances. However, recent studies showed that the genetic variation is already high on small geographic scales, yet direct comparisons of the genetic variation across different geographic scales are rare. Here, we present a new local diversity panel covering 19 km^2^ with accessions of the Veluwe, the Netherlands. We compared the genetic variation of this local diversity panel to a national and a continental *A. thaliana* diversity panel. Direct comparison of these three geographic scales showed that local accessions harbour already 41.8% of the genetic variation found on a continental scale despite the substantial difference in geographic surface area covered. Moreover, a rapidly ascending logarithmic relationship between genetic and geographic distances was observed at continental, national and local scale and thus irrespective of the geographic scale considered. The high level of local genetic variation reported here poses new questions on which evolutionary forces are driving and maintaining this, and how much this constrains experimental design when using local *A. thaliana* populations in future eco-evolutionary studies.

## Introduction

Over the past decades, the plant species *Arabidopsis thaliana* became a model for molecular genetics. Currently, *A. thaliana* is increasingly used in ecological and evolutionary studies, which harness the vast amount of (genetic) resources and knowledge that have been established (1–3). These developments have led to the establishment of new regional and local *A. thaliana* diversity panels (4–6). Such diversity panels can be used to link evolutionary and adaptive processes to particular environmental contexts, and search for specific mechanisms or genes that may underly adaptation (3). The use of *A. thaliana* for eco-evolutionary studies depends on two important features. First, *A. thaliana* spans immense native range across Eurasia, including a wide variety of environments (7). Second, *A. thaliana* displays considerable genetic variation (8–10).

The amount and structure of the genetic variation in *A. thaliana* across its native range have been studied on various scales and at different locations (9–13). On a continental scale, it was discovered that *A. thaliana* originated from Africa, from where it spread to its current native range using refugia such as the Iberian Peninsula during glacial periods. In these refugia distinct and conserved ‘relict’ genetic variation was found, while the rest of the more recently populated Europe harbours a mixture of genetic variation with lower genetic relatedness (14,15). This genetic variation has a geographic distribution with a strong longitudinal gradient on a continental scale (10,12–14,16,17). A positive correlation between the genetic and geographic distances was found and postulated as the genetic model of ‘isolation by distance’ (IBD) of which the strength and type (i.e. shape) can vary depending on the geographic region or scale (13,14). IBD is driven by evolutionary processes such as genetic drift, local adaptation, dispersal limitations, as well as plant-related traits, such as self-fertilization (18,19). Inherently, the IBD model suggests that the smaller the geographic distance, the greater the genetic similarity between accessions (i.e. smaller genetic distances). However, more recently it was argued that the relatively simple IBD model does not conform to the real distribution of *A. thaliana* genetic diversity (12).

Additional studies of populations revealed that *A. thaliana* harbours more genetic variation on smaller geographic scales than would be expected based on IBD from a self-fertilizing plant species with an outcrossing rate of 1 to 3% (20). This was shown on a regional scale by various studies across Europe such as France, Sweden, and the Iberian Peninsula (9,21,22). Moreover, examination of local diversity panels in Germany, Spain and France showed substantial variation in genetic diversity and outcrossing rates where even closely located sites displayed strong genetic differentiation (4,5,23,24). Yet, direct comparisons between the genetic variation across different geographic scales are limited. Using simulations with Eurasian accessions, Platt et al. (2010) already discovered that the relationship between genetic and geographic distances is similar on different geographic scales. Recently, Schmitz et al. (2023) showed that within local urban populations similar genetic distances were found as among accessions across Europe. As urban populations contain less genetic variation than rural populations, the question is how local rural populations compare to larger geographic scales (23).

The aim of the present study was to relate local genetic variation in a habitat with modest diversity in ecological niches (i.e. climate, altitude and soil composition), to larger geographic scales. To investigate this, we introduce a new local diversity panel with accessions from small area (19 km^2^) of the Veluwe, a region located in the central-east of the Netherlands. This area is largely rural with nature reserves intermingled with small-scale (former) agricultural fields and is entirely situated on one single parent soil material with relatively few climatic differences (25). To put this local genetic variation in perspective of larger geographic scales, we compared this local panel to a national panel completely covering the Netherlands (26) and a subset of a European continental diversity panel (12). We compared the genetic variation of local to continental diversity panels by joint variant calling, revealing the existence of surprisingly high amounts of genetic variation on a local scale with same climate and similar soil composition. We also showed that genetic distances increase by a steep logarithmic function that rapidly grows over short geographic distances.

## Results

### Experimental design of the study

The focal point of our local-to-continental analysis of the genetic variation is a set of 51 *A. thaliana* accessions from agricultural and abandoned agricultural fields, as well as road verges, within 19 km^2^ of the Veluwe, Netherlands (25). In addition, we used the DartMap dataset with 192 accessions that are evenly sampled across the Netherlands (26) and a subset of 230 accessions of the 1001G dataset, collected within a range of 0-1000 km away from the Veluwe (12). In order to compare the genetic variation at different geographic scales, we subjected the three datasets to a joint variant calling pipeline based on whole genome short read re-sequencing (Illumina). We identified bi-allelic SNPs and small insertions and deletions (collectively referred to as genetic variants) in comparison to the *A. thaliana* Col-0 reference genome.

First, we validated the integration of the three datasets by analysing the population structure in the organellar and nuclear genomes. We expected that, irrespective of the dataset of origin, the organellar genome would show clusters according to ancient genetic variation and that the nuclear genome would show a geographic pattern due to more recently established genetic variation (5,8,27). Next, we set out to compare the different geographic scales and hereby studied the overlapping genetic variants and the genetic versus geographic distances across the three geographic scales. To compare the geographic scales, we had to account for differences in sample size and sampling density between the three datasets. Therefore, we nested the accessions based on their geographic scale; local, national or continental (respectively referred to as Veluwe Plus, Netherlands and Europe). In this way, local genetic variation is part of the national genetic variation, and the national genetic variation is part of the continental genetic variation. To assess the geographic distances, we determined pairwise distances between all plant sampling coordinates. To assess the genetic distances, we determined the number of genetic variants different between accessions. An overview of the location of the accessions across the different geographic scales is depicted in **Figure 1**.

**Figure 1.**
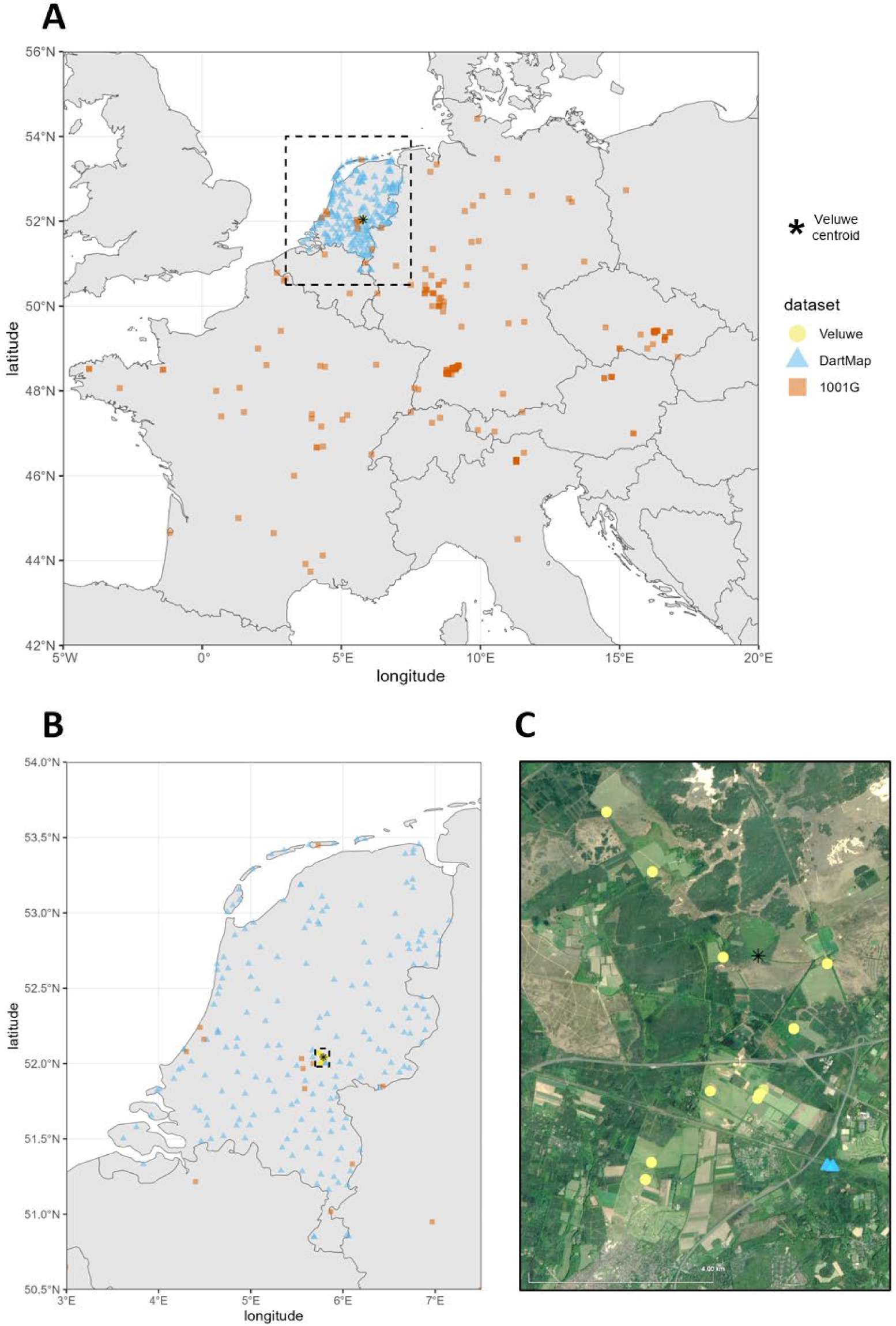
Locations of the *A. thaliana* accessions at different geographic scales. (**A**) Continental scale (Europe) with all 473 accessions, dashed rectangular indicates location of the Netherlands. (**B**) National scale (Netherlands) with 253 accessions, dashed rectangular indicates location of the Veluwe. (**C**) Local scale (Veluwe Plus) with 53 accessions. Black asterisk indicates Veluwe centroid, colours and shapes indicate dataset.

### Population structure of organellar and nuclear genetic variation validates integration of datasets

We identified 277, 629 and 975 genetic variants in the chloroplast genome and 160, 252 and 494 genetic variants in the mitochondrial genome in the Veluwe, DartMap and 1001G dataset, respectively. Assessing the organellar population structure revealed three distinct clusters in both the chloroplast (**Fig. S1A**) and mitochondrial (**Fig. S1B**) genome. The three chloroplast clusters are nearly similar in composition to the three mitochondrial clusters except for 17 out of 473 accessions (**Table S1** and **S2**). This suggests that the majority of accessions belong to a single ancestral cluster. Each cluster contains accessions from each dataset and so all three datasets have accessions from all three ancestral clusters (**Table S1** and **S2**). In conclusion, these results showed that dataset driven patterns (e.g. population structure according to dataset) are absent and supports the integration of the three datasets.

Through our analysis of the nuclear genome, we identified 2,464,107 genetic variants in the Veluwe dataset, 4,176,425 genetic variants in the DartMap dataset and 5,144,498 genetic variants in the 1001G dataset. The nuclear genetic variation has a strong geographic pattern, distinguishing clusters originating from different countries and separating western European countries from eastern European countries along PC1 (**Fig. S2A**). The Veluwe dataset completely co-localized with the DartMap dataset while the 1001G dataset only partly overlapped (**Fig. S2B**). This indicates that the Veluwe accessions at 19 km^2^ share the genetic variation found in the accessions within the entire Netherlands (40,040 km^2^). This is confirmed by separately analysing all accessions originating from the Netherlands (n = 253) where all three datasets co-localized (**Fig. S2C**). To complement the previous findings, we also estimated the ancestry of the accessions using admixture analysis and detected again a geographic pattern. With increasing ancestral clusters (K = 3, K = 4 and K = 6) a Dutch (orange), French (green), south-east European (blue), Italian (grey), and German (pink) dominated ancestry could be distinguished (**Fig. S3**). Compared to other accessions from the Netherlands, the Veluwe accessions had relatively lower mixed ancestry and certain ancestral lineages were absent from the Veluwe dataset, while they were present in other accessions from the Netherlands (**Fig. S3**). This corroborates that the local genetic variation of the Veluwe accessions forms smaller part of the genetic variation present in the Netherlands. Most importantly, the observed geographic structure in the nuclear genetic variation in our results is in accordance with the previously identified pattern (5,8). Thereby, dataset-driven patterns are absent which validated the integration of the three datasets.

### Local accessions harbour a considerable part of the genetic variation found on a continental scale

To understand how the level of genetic variation compares at the three different geographic scales, we assessed the amount of shared and unique nuclear genetic variants. Between the datasets, there was a core set of 36.9% of the genetic variants (**Fig. 2A**). The Veluwe dataset harboured few unique genetic variants (1.15%) sharing the majority of its variants (98.85%) with the DartMap and 1001G dataset. This finding is consistent with our observation that Veluwe genetic variation falls within the genetic variation of the Netherlands and Europe (**Fig. S2 BC**). The 1001G dataset had more unique genetic variants (27.7%) compared to the DartMap (11.1%) even though both datasets had a comparable number of accessions (respectively 230 and 192) which can be explained by the increased geographic area covered by the 1001G accessions. Next, we addressed how many genetic variants were shared between the different geographic scales. The increase from local to national to continental scale are large geographic increments in the surface covered but strikingly, the number of genetic variants in the nested geographic scales increased less steeply (respectively, an approximate 52,000 times increase in geographic area compared to a 2.4 times increase of the number of genetic variants, **Table S3**). The local accessions already contained 41.8% of all genetic variants present in all accessions at the continental scale (**Table S3** and **Fig. 2B**). Going from local scale to national scale recovered an additional 31.8% of genetic variation so that it encompassed 73.6% of all genetic variants present at the continental scale (**Fig. 2B**).

**Figure 2.**
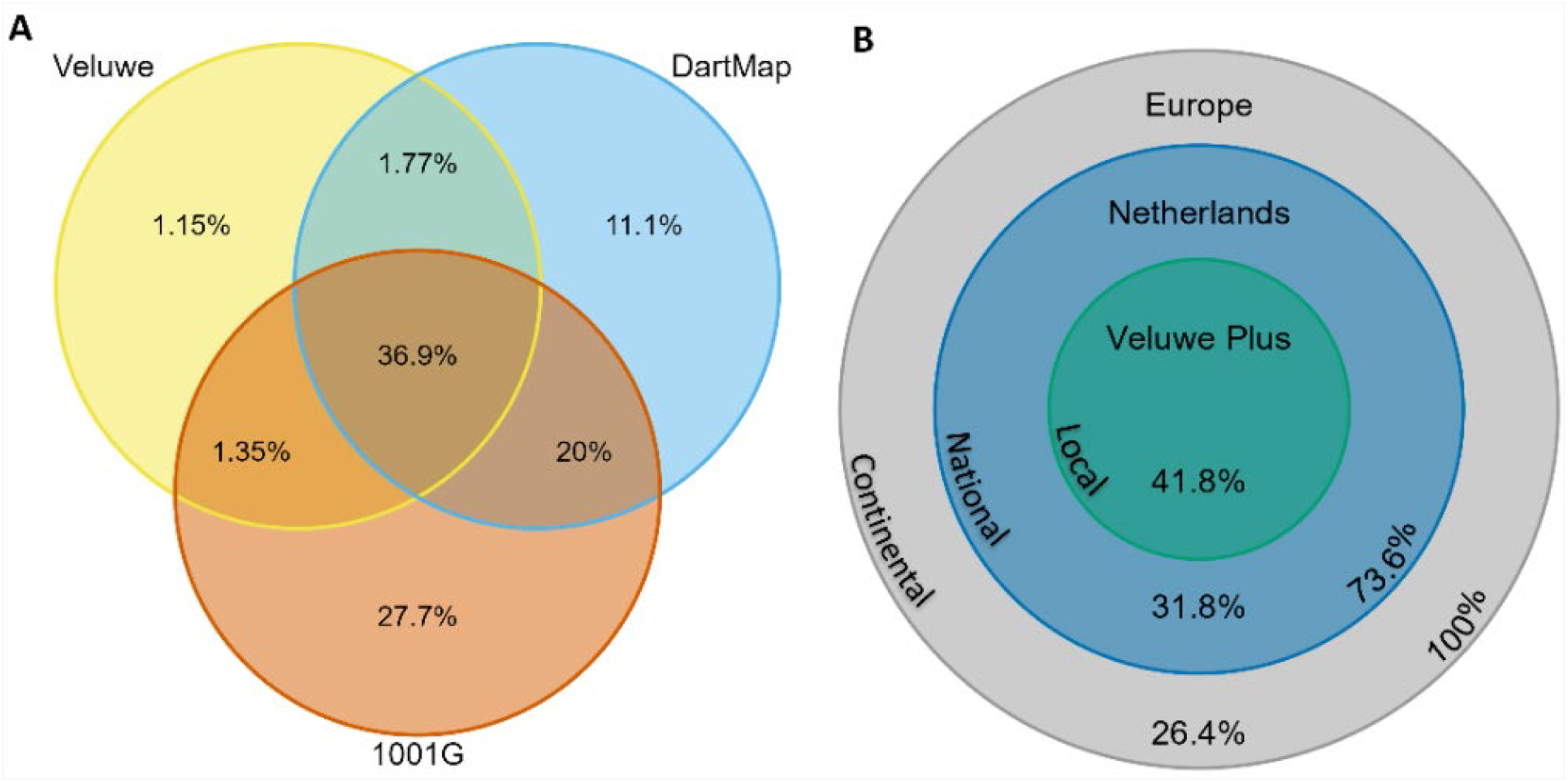
Venn diagrams showing overlapping and unique genetic variants in percentages. (**A**) Based on dataset (Veluwe n = 51, DartMap n = 192, 1001G n = 230). Colour indicates dataset. (**B**) Based on geographic scale (Local Veluwe Plus n = 53, National Netherlands n = 253, Continental Europe n = 473). Colour indicates geographic scales. Percentages on border of circles indicate cumulative of inner circle percentages.

### Steep logarithmic increase of genetic distance across geographic scale

Lastly, we compared the pairwise genetic and geographic distances between accessions across different geographic scales. At all geographic scales there was a significant correlation between genetic and geographic distances. However, this correlation was weaker at local (Mantel *r* = 0.20, *p* = 0.001) and national (Mantel *r* = 0.22, *p* = 0.001) scales than at the continental scale (Mantel *r* = 0.50, *p* = 0.001). This suggests that at smaller geographic scales, an increase of geographic distance correlates less to an increase in genetic distance. When plotting the genetic versus geographic distances, there was a highly similar relationship for all geographic scales (**Fig. 3A**, **B** and **C**). The continental, national and local all showed a logarithmic relationship (all three *p* < 0.001) with initial strong increase of genetic variants over the shortest geographic distances which plateaus rapidly around 1.25 million genetic variants (respectively **Fig. 3A**, **B** and **C**). Since the logarithmic models of the national and continental scale have nested data, we also compared the logarithmic models per dataset. Here, we again saw the same pattern that with all geographic distance studied (up to 1500 km, 300 km or 10 km) the logarithmic models describing the genetic versus geographic distance were similar for all datasets (**Fig. S4**). The only exception was the DartMap dataset, which had a less steep increase of genetic distance at the first 5 km of geographic distance (**Fig. S4C**). This is most likely due to the relatively small number of accession pairs in the DartMap dataset at such small geographic distances. Lastly, we compared solely the genetic distances in the different datasets (**Fig. 3D**). The mean pairwise genetic distance between accessions in the Veluwe dataset (1,116,478 genetic variants) was only slightly lower than the mean genetic distances in the DartMap and 1001G dataset (respectively 1,205,739 and 1,213,625 genetic variants). All in all, these results corroborated that the genetic distances between accessions were comparable between datasets and the curves describing the genetic versus geographic distances are similar, irrespective of when considering a continental, national or local scale.

**Figure 3.**
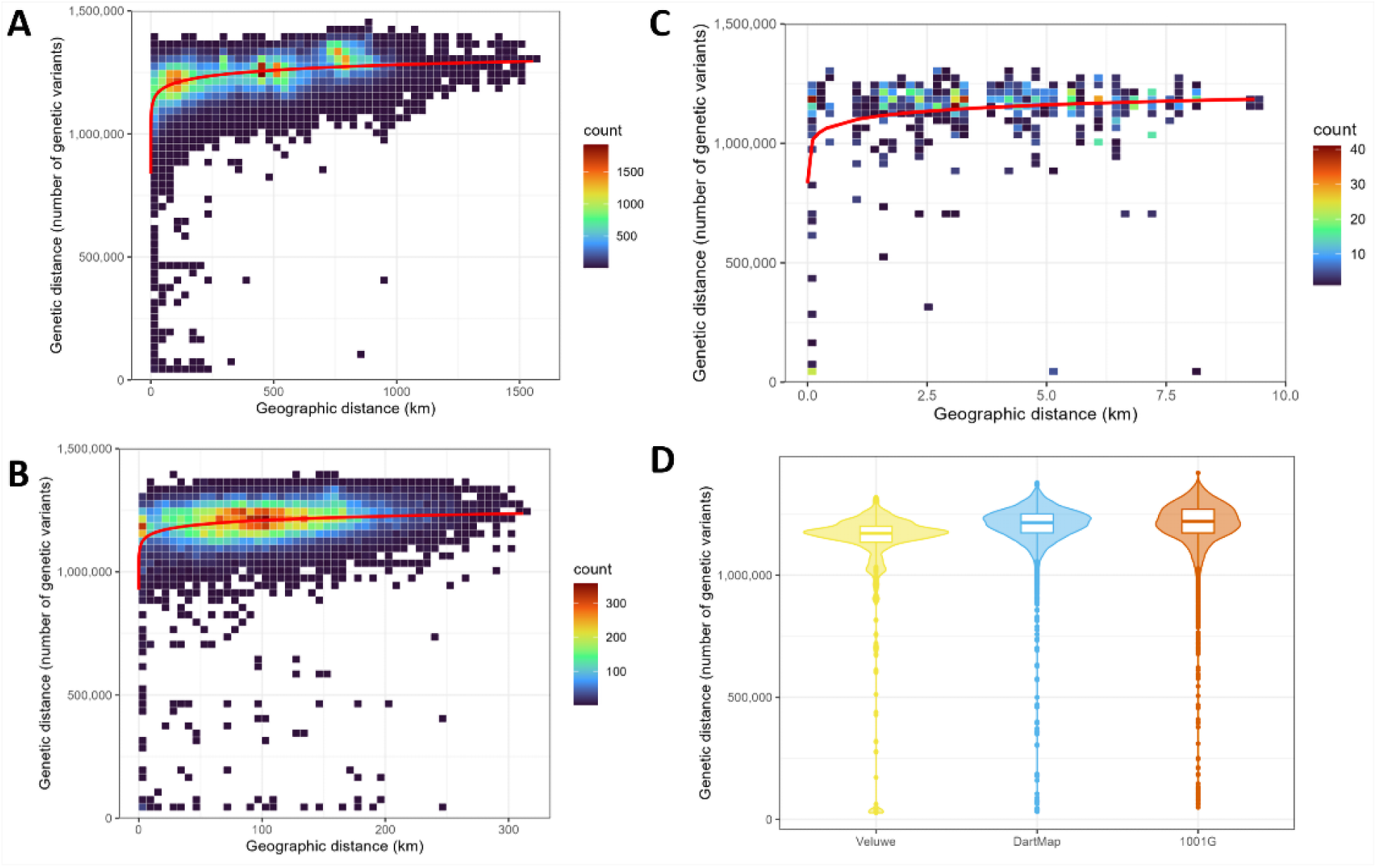
Genetic versus geographic distances across different geographic scales. (**A**) Scatter bin plot of genetic versus geographic distance in Europe, (continental scale n = 473). (**B**) Scatter bin plot of genetic versus geographic distance in the Netherlands (national scale n = 253). (**C**) Scatter bin plot of genetic versus geographic distance in Veluwe Plus (local scale n = 53). Red lines indicate fitted logarithmic model for each geographic scale. X-axis displays pairwise geographic distance between accessions, expressed in kilometre. Y-axis displays pairwise genetic distance between accessions, expressed as number of genetic variants. (**D**) Pairwise genetic distances per dataset in boxplot including density. Genetic distance is expressed as number of genetic variants different in a pairwise comparison between accessions.

## Discussion

We studied how genetic variation at a local scale relates to larger geographic scales and introduced a novel diversity panel of *A. thaliana* accessions from a small geographic area with a relatively uniform soil type and climate. We integrated three different datasets to be able to accurately compare local, national and continental scale and validated this by studying the organellar and nuclear population structure. Comparing the chloroplast and mitochondrial genome revealed three distinct organellar clusters, which is in line with the three highly diverged haplogroups representing ancient variation predating post-glacial expansion (26,27) (**Fig. S1**). In the nuclear population structure, we observed a geographic pattern at the European scale with a longitudinal gradient and clustering of accessions from different countries (**Fig. S2** and **S3**), thought to represent more recent genetic variation following the last glacial expansion (5,8,11). Within this pattern, the Veluwe panel was nested within the genetic variation of the Netherlands (**Fig. S2BC**).

The Veluwe panel forms a concentrated sampling area and it revealed a high amount of genetic variation. On all three geographic scales, we observed a correlation between genetic and geographic distances, which was most strongly displayed on a continental scale (Mantel *r* = 0.50) compared to national (Mantel *r* = 0.22) and local (Mantel *r* = 0.20) scale. However, the logarithmic relationship between the genetic and geographic distances turned out to be highly comparable when considered at local, national and continental scale (**Fig. 3**). Thereby, our results show isolation by distance but this seems to be independent of geographic scale. This confirms the previously simulated work presented by Platt et al. (2010) as well as the previous finding that IBD among non-relict accessions only occurs at very small distances (12). Moreover, our findings are in line with the recent results that local urban *A. thaliana* accessions show genetic distances of the same order as on a continental scale (5). All in all, this leads to the conclusion that the genetic variation (SNPs and indels) at a geographic distance of 10 km can be as much as over a geographic distance of 1000 km.

Despite the fact that the Veluwe area spans just 19 km^2^, these 53 local accessions displayed 41.8% of the genetic variants detected in 473 accessions within 1000 km radius of the Veluwe (**Fig. 2B**). Only a small fraction (1.15%) of these variants are unique to the Veluwe dataset, suggesting that genetic variation is mainly composed of variants that are distributed across larger geographic regions (**Fig. 2A**). This observation is not unique to the Veluwe, as urban accessions from Cologne also form a subgroup from the genetic variation present in Germany and surrounding countries (5). This suggests that local hotspots of high genetic diversity are not unique and actually present in both rural and urban systems. However, high genetic diversity on a local scale is not a given, as populations with very little genetic variation or genetic diversity are also reported and for example make up to 24% of the studied populations of the Iberian Peninsula (6,23,28,29). There is ample evidence that environmental factors have the ability to shape the genetic variation in a plant population to optimize plant performance (11,30–33). In addition, there are ongoing evolutionary processes to consider, such as drift and gene flow, which may confound with environmental factors (6). Decreasing sequencing costs and thereby increasing sequencing efforts of new (local) populations will help to illuminate why certain populations display particularly low or high levels of genetic variation.

As a self-fertilizing species, *A. thaliana* is generally presumed to rarely outcross, having an average outcrossing rate of 1-3%. However, large variation in the outcrossing rate has been observed, going up to 14% (23,29). Although heterozygosity appears to be constrained by a low outcrossing rate, it is sufficient to generate and distribute a considerable amount of genetic diversity at a local scale (20,23). At present, we cannot discern what factors might have contributed to the observed genetic variation in the Veluwe but there could be various reasons. First, there may have been human-mediated gene flow (14). As the sampled Veluwe sites have experienced human management, we could foresee a potential effect of such human-intervention. Second, rural environments display greater genetic diversity compared to urban sites, which fit the description of the sampled Veluwe area (23). Third, the temporal aspect of local genetic variation is increasingly acknowledged. Standing genetic variation is known to change over different years as immigration or germination from seed banks can provide new gene flow (23,24). Future studies may give insights in how locally these high levels of genetic variation arise and what evolutionary processes might maintain them.

We highlighted the remarkable amount of genetic variation present locally in the Veluwe. Numerous studies have already demonstrated the value of smaller-scale diversity panels to complement the established continental and global panels (21,22,31). Even genetic variation of an *A. thaliana* population within 1 km was proven insightful for the identification of novel defensive metabolite genes (34). Similar to this, the Veluwe diversity panel can serve as a starting point for future research on local adaptation, as this region is characterized by differences in abiotic and biotic soil composition in the absence of climatic clines (25). However, the presence of complex genetic variation at a local scale also presents challenges to make meaningful correlations with environmental factors or the detection of causal genes that contribute to adaptation. Therefore, future studies using new local diversity panels should adjust their sampling strategy and analyses accordingly.

## Materials and methods

### Sampling and plant propagation

*Arabidopsis thaliana* is a colonizing annual plant of disturbed rural soils, but also able to successfully establish in urban environments (1). In May and June 2018 at the end of their flowering season, 51 dry seed-carrying *A. thaliana* plants were collected from eleven different sites in the South Veluwe region in the Netherlands (**Fig. 1C**). These sites cover an area of 19 km^2^ and consist of the same parent soil material and have the same climatic conditions (25). The eleven sites consist of two agricultural fields, two grassy road verges and seven former agricultural fields turned into natural grasslands and are considered (semi) rural. At each site, two to six plants were collected from larger standing populations and plants of the same site share the site coordinates, indicated in **Table S4**. The mean geographic distance between sites is 3.7 km with a maximum of 8.2 km and a minimum of 56 meters between sites. The generation 1 (G1) seeds were harvested from the field-sampled plants and propagated to generation 3 (G3) plants through single-seed-descent by self-fertilization in a climate chamber (with 16-hour light at 23°C and 8-hour dark at 21°C cycles).

### DNA isolation and sequencing

Pooled seedling tissue and leaf tissue was used for DNA isolation. Plant material was frozen using liquid nitrogen and ground to fine dust. Material was incubated in 500 µl preheated (65°C) CTAB buffer (2% CTAB, 1.4 M NaCl, 100 mM Tris-HCl, 20 mM EDTA, pH 8) for 60 min at 65°C. 500 µl of chloroform was added, mixed and centrifuged (14000 rpm for 10 min) and supernatant was collected. 500 µl of ice-cold isopropanol was added, mixed and centrifuged (14000 rpm for 10 min) to precipitate the DNA. The precipitate was washed with 70% ethanol and air dried after which the DNA was dissolved in TE buffer (10 mM Tris-HCl, 1 mM EDTA, pH 8). The dissolved DNA was treated with RNase A (Sigma-Aldrich) for 45 min at 37°C after which the same precipitation, washing and dissolving steps were repeated. Genomic DNA was sent out to Novogene (Cambridge, United Kingdom) for library preparation and sequencing of paired-end 150-bp reads using Illumina NovaSeq 6000 aiming for at least 30x coverage.

### Sequence data collection

Raw sequence data of the 51 accessions in fastqc format (European Nucleotide Archive: PRJEB63356) are referred to as the ‘Veluwe’ dataset and was used as a local scale. For the national scale, the raw sequencing data of the DartMap panel consisting of 192 Dutch accessions was used, referred to as the ‘DartMap’ dataset (Sequence Read Archive ID: PRJNA727738) (26). A subset from the 1001G Project (Sequence Read Archive ID: SRP056687) was used as a continental scale, which is referred to as the ‘1001G’ dataset (12). For this subset, we selected accessions with coordinates within 1000 km of the Veluwe (Veluwe centroid; longitude 5.786972 and latitude 52.04302) and excluded accessions from locations that had a body of water separating them from the Veluwe (accessions from Finland, Ireland, Norway, Sweden, United Kingdom, and the islands of Denmark). There were a small number of accessions that were sequenced with Illumina Genome Analyzer II or with an unknown sequencing machine. These accessions had either very short (< 50 bp) or very long (> 250 bp) reads and the reverse and forward reads were not indicated. The length and/or lack of indication of read direction might be a confounder and therefore these accessions were excluded. We referred to the remaining 252 accessions as the 1001G dataset. These three datasets yielded a total of 495 accessions used as input in the variant calling pipeline.

### Variant calling pipeline

Genetic variants were either single-nucleotide polymorphisms (SNPs) or short insertions or deletions. These were called in using a workflow adapted from (26) and based on the Genome Analysis Toolkit (GATK) Best Practices (35). GATK toolkit v4.3.0.0 was used for various steps of the pipeline (36). For all steps default settings were used, unless specified differently. Due to poor sequence quality, 22 accessions of the 1001G dataset were removed in the variant calling pipeline. The output of the variant calling pipeline consisted of six VCF files, each with 473 accessions. There were two nuclear, two chloroplast and two mitochondrial VCF files with hard-filtered biallelic variants. Per genome, one VCF file was without MAF filtering and the other VCF file with MAF filtering (0.10 for nuclear genome and 0.05 for chloroplast and mitochondrial genome). The VCF files were used in subsequent data analysis. A detailed description of the variant calling pipeline can be found in the supporting information (**Text S1**)

## Data analysis

### Data structure

For the data analysis we distinguished two ways of structuring the 473 accessions; based on dataset or based on geographic scale. For comparing between the datasets, the accessions were grouped on the basis of the original diversity panel or dataset; ‘Veluwe’ with 51 accessions, ‘DartMap’ with 192 accessions or ‘1001G’ with 230 accessions. For comparing geographic scales, the accessions were grouped using nesting on the basis of their geography; local scale ‘Veluwe Plus’ with 53 accessions (51 of Veluwe and 2 of DartMap), national scale ‘Netherlands’ with 253 accessions (51 of Veluwe, 192 of DartMap, and 10 of 1001G) and continental scale ‘Europe’ with 473 accessions (51 of Veluwe, 192 of DartMap, 230 of 1001G). The spread of the accessions across the different geographic scales is visualized using packages rnaturalearth (v0.3.2), rnaturalearthhires (v0.1.0) and Google Earth Pro (**Fig. 1**). For part of the data analyses, we used R (v4.2.1) and RStudio (v.2022.12.0) with various packages (37). Results were visualized using RStudio and package ggplot2 (v3.4.0).

### Number of genetic variants

For determining the number of genetic variants, we used the VCF without MAF filtering to also take rare variants into account. The genetic variants were determined separately for the nuclear, chloroplast and mitochondrial genomes. To determine the number of genetic variants for each dataset (either Veluwe, DartMap or 1001G) we made subsets of the relevant accessions using bcftools view (v2.5.2). Next, we counted the variants which were still variable within the subset of accessions using GATK SelectVariants (v4.3.0.0) and CountVariants.

### Population structure

For the population structure analysis, we used the VCF with MAF filtering to prevent an excessive effect of rare variants on the population structure and analysed this on basis of dataset structure. Package SNPRelate (v1.32.3) was used to perform principal component analysis (PCA) on the nuclear, chloroplast and mitochondrial genome separately. The first two principal components were plotted to visualize the population structure. Chloroplast and mitochondrial PCAs were coloured by dataset. We performed hierarchical clustering with complete linkage as agglomeration method on PC1 and PC2. From this, three clusters were determined using package stats (v4.2.1), arbitrarily named A, B or C for further analysis. Cluster composition comparison between the chloroplast and mitochondrial genome was done manually. The nuclear PCA was coloured by country and by dataset and included 0.95 confidence interval ellipses. In addition to the nuclear PCA, we also ran ADMIXTURE (v1.3.0) for ancestry estimation based on the nuclear genome which applies a maximum likelihood estimation specific for multi-locus SNP genotype datasets. For this analysis we used the VCF without MAF filtering as ADMIXTURE addresses variant frequency/occurrence inherently. To run ADMIXTURE, we used plink (v1.90b6.21) to transform the VCF to a bed file (38). We ran ADMIXTURE for number of clusters K = 1 to 10 and excluded Ks with high cross-validation errors (K = 1, 2, 5) for further analysis.

### Overlapping genetic variants

For determining the number of overlapping genetic variants, we used the VCF without MAF filtering to also take rare variants into account and analysed this both on the basis of dataset structure and geographic scale structure. Next, we assessed the number of overlapping genetic variation between the datasets or between geographic scales. This was only determined for the nuclear genome. We made subsets of the relevant accessions (per dataset or geographic scale) using bcftools view (v2.5.2). Next, we selected the variants which were still variable within the subset of accessions using GATK SelectVariants (v4.3.0.0) and those variants were submitted to bcftools isec to calculate the number of overlapping variants between the subsets. Package VennDiagram (v1.7.3) was used for visualization.

### Genetic versus geographic distance

The genetic distance was used to determine the genetic (dis)similarity between accessions and measured as Hamming distance which equals the number of genetic variants different between accessions. For calculating the genetic distances, we used the VCF without MAF filtering to also take rare variants into account and analysed this both on basis of dataset structure and geographic scale structure. The relationship between genetic and geographic distances were only analysed for the nuclear genome and all analyses were conducted in RStudio. Genetic variants were first transformed to a binary system with 1s for homozygous variants and 0s for homozygous reference and heterozygous variants. Based on this binary system a genetic distance matrix was determined in Hamming distances. For these steps we used packages vcfR (v1.14.0) and poppr (v2.9.4). Geographic distances between accessions were computed using package sp (v1.6-0). The genetic and geographic distance matrices contained all pairwise comparisons between all accessions. We performed a Mantel test with Pearson correlation and 999 permutations comparing the genetic and geographic distances matrices using package vegan (v2.6-4). We fitted a logarithmic model (genetic distance ∼ log(geographic distance) using package stats (v4.2.1) and used the logarithmic model to calculate predicted values for in the plots. This has been done both per geographic scale and per dataset.

## Additional information

## Data availability

Raw sequence data of the 51 accessions in fastqc format is available on the European Nucleotide Archive (PRJEB63356). The DartMap raw sequence data of 192 accessions is available on the European Nucleotide Archive (PRJNA727738). The 1001G Project raw sequence data is available through the Sequence Read Archive of NCBI (SRP056687).

## Author contributions

V.H, B.S, V.W and W.vdP developed the study concept and design; V.H, V.W and W.vdP collected the Veluwe A. thaliana accessions; V.H propagated the Veluwe A. thaliana accessions from G1 to G3; V.H performed the DNA isolations; M.G.M.A provided the DartMap dataset; R.W. designed the variant calling pipeline; Y.B. made variant calling pipeline and processed the raw sequencing data; V.H and Y.B performed the data analysis; R.B, R.W. and R.H provided input on the variant calling pipeline and data analyses; V.H, B.S, V.W. and W.vdP drafted the manuscript; M.G.M.A, R.B., R.W. and Y.B. provided input on the manuscript. All authors read and approved the final manuscript.

## Acknowledgements

We thank Jeanne Brives for her help with the DNA isolations and Koen Verhoeven for the insightful suggestions and comments on the manuscript.

## Supporting information

**Text S1** | Detailed description of variant calling pipeline and filtering

In first part of the pipeline, the separate raw sequencing files were processed, quality checked and mapped to the reference genome per accession. First, the *A. thaliana* reference genome (TAIR10, European Nucleotide Accession number: GCA_000001735.2, accessed via https://www.ebi.ac.uk/ena/browser/view/GCA_000001735.2?show=blobtoolkit) was indexed using bwa index (v0.7.1.17-r1188) and samtools faidx (1.16.1) and a GATK directory was created using GATK CreateSequenceDictionary. Raw sequencing reads were quality-trimmed based on their Phred score using CutAdapt (v4.1) (39). When the Phred score was lower than 20 (on a 33 offsett Illumina > 1.8 scoring system), the 5’ and 3’ ends were trimmed and if less than 2/3 of the original read length remained, the read was filtered out. CutAdapt ran separately for Veluwe and DartMap accessions (raw read length around 150 bp, cut-off 101 bp) and 1001G accessions (raw read length around 100 bp, cut-off 67 bp) (settings: -q 20, -m 67 or 101, and --pair-filter any). Before (raw reads) and after (trimmed reads) CutAdapt, the reads were submitted to FastQC (v0.11.9) to check for aberrant quality levels. The trimmed reads were mapped to the *A. thaliana* reference genome using bwa mem (v0.7.17-r1188) (40). In bwa mem, read groups were specified for accessions from Veluwe and DartMap datasets but read group information (including sequencing machine, lane and cell) was unavailable for 1001G dataset. To prevent interferences later in base quality score recalibration unique read groups were created using GATK AddOrReplaceReadGroups and therefore, sequencing information was not accounted for in the 1001G accessions. The resulting alignment files were sorted, compressed and indexed using samtools (v 1.16.1) (41).

In second part of the pipeline, certain accessions were filtered out and genetic variants were called. Using GATK Picard MarkDuplicates the alignment files of different libraries from the same accession were merged and duplicate read pairs were marked (settings: -- OPTICAL_DUPLICATE_PIXAL_DISTANCE 2500). Using samtools index, the merged bamfile was indexed. GATK CollectAlignmentSummaryMetrics, bamtools stats, and samtools coverage were used to check genome coverage and sequencing depth. Percentage reads mapped and sequencing depth of the nuclear genome is indicated for each dataset in **Figure S5**. We defined three selection criteria for the accessions; no aberrant FastQC score after trimming, minimal sequencing depth after alignment to reference genome of 10x, and minimum of 80% of reads mapped. In total 22 accessions (all from 1001G dataset) were filtered on the basis of the selection criteria, resulting in 473 samples fit for variant calling. Using GATK BaseRecalibrator and GATK ApplyBQSR the aligned bases were recalibrated (settings: as --known-sites) using a reference SNP set called in 1135 *A. thaliana* accessions of 1001G consortium (release 37a, accessed via http://ftp.ensemblgenomes.org/pub/plants/release-37/vcf/arabidopsis_thaliana/) (12). The resulting bam file was indexed using samtools index. Genetic variants compared to the reference genome were called in each sample using GATK HaplotypeCaller. This was done in -ERC GVCF mode and a maximum of 3 alternate alleles at each site was allowed. Accessions were genotyped using GATKGenomicsDBImport and GATK GenotypeGVCFs. Three VCF files were generated for respectively the nuclear, chloroplast and mitochondrial genome.

In the third and last part of the pipeline, genetic variants were filtered separately for each VCF to remove false positive calls. Nuclear variants were recalibrated based on the reference SNP set called in 1135 *A. thaliana* accessions of 1001G consortium (release 37a) (12). GATK VariantRecalibrator recalibrated called nuclear variants separately in SNP more and indel mode (settings: --trust-all-polymorphic -tranche 99.9 -an QD -an MQ -an MQRankSum -an ReadPosRankSum -an FS -an SOR -an DP -an ExcessHet -an InbreedingCoeff --max-gaussians 6 for SNP mode or --max-gaussians 4 for indel mode). The reference SNP set was used as training and truth set during recalibration (settings: --prior=10.0). GATK ApplyVQSR was used to apply the information of GATK VariantRecalibrator, separately in SNP mode and indel mode (settings: --truth-sensitivity-filter-level set 99.9). Nuclear, chloroplast and mitochondrial variants were additionally hard-filtered using GATK VariantFiltration and a set of thresholds determined by (26) (settings: nuclear QD < 21.0, FS > 70.0, SOR > 3.0, MQ < 30.0; mitochondrial QD < 2.0, FS > 100.0, SOR > 4.0, MQ < 20.0; chloroplast QD < 22.0, FS > 70.0, SOR > 3.0, MQ < 30.0).

Hard-filtered variants were removed using vcftools (v0.1.16) and after GATK CollectVariantCallingMetrics was used to perform a quality control (settings: --DBSNP with reference set). We assessed the number of hard-filtered genetic variants per genome and saw that although the number of variants filtered varied per dataset, the distribution of remaining genetic variants was even between datasets (**Fig. S6**). Next, multiallelic variants were filtered using vcftools to only retain the biallelic variants, as these variants are called with a higher confidence. The last step of variant filtering was the application of a minor allele frequency (MAF) and removal of rare variants, again using vcftools. We used a MAF of 0.10 for the nuclear genome and 0.05 for the chloroplast and mitochondrial genomes. A lower MAF for the organellar genomes was used to prevent discarding a relatively large portion of variants. At the end of the variant calling pipeline there were two nuclear, two chloroplast and two mitochondrial VCF files with hard-filtered biallelic variants (without and with MAF filtering) for a total of 473 accessions.

**Table S1.** Overview of the number of accessions in the chloroplast genome clusters. Number of accessions is indicated per dataset and totals per cluster and per dataset are given.

| dataset | cluster A | cluster B | cluster C | total |
| --- | --- | --- | --- | --- |
| Veluwe | 29 | 3 | 19 | 51 |
| DartMap | 62 | 52 | 78 | 192 |
| 1001G | 48 | 107 | 75 | 230 |
| total | 139 | 162 | 172 | 473 |

**Table S2.** Overview of the number of accessions in the mitochondrial genome clusters. Number of accessions is indicated per dataset and totals per cluster and per dataset are given.

| dataset | cluster A | cluster B | cluster C | total |
| --- | --- | --- | --- | --- |
| Veluwe | 30 | 2 | 19 | 51 |
| DartMap | 75 | 39 | 78 | 192 |
| 1001G | 51 | 104 | 75 | 230 |
| total | 156 | 145 | 172 | 473 |

**Table S3.**
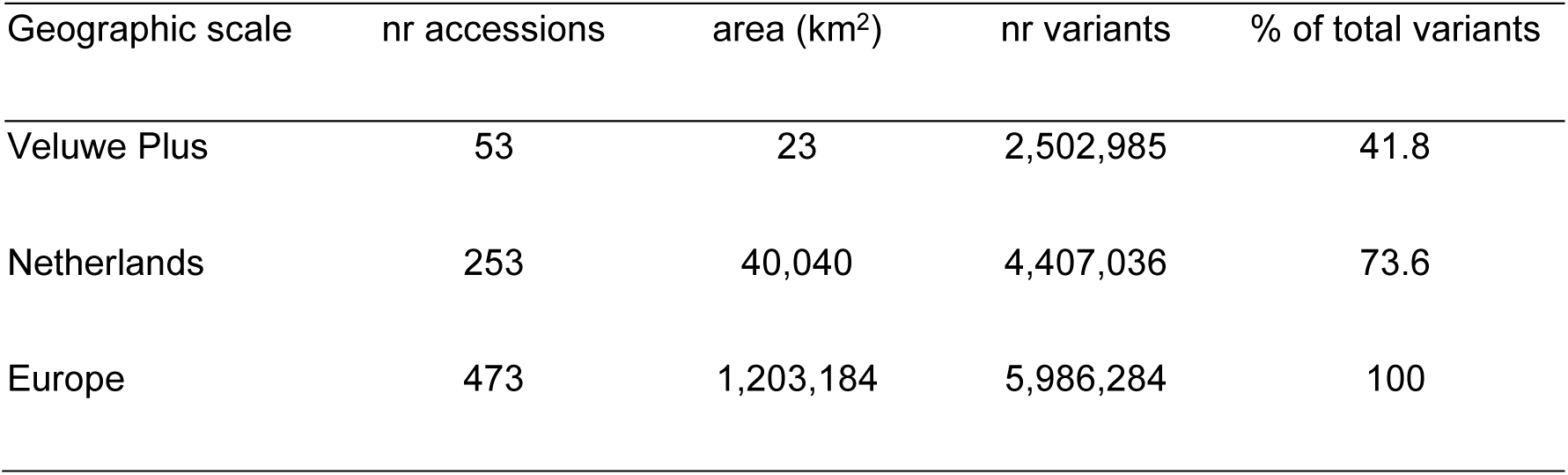
Number of genetic variants per geographic scale. Number of accessions, area covered by the accessions, number of genetic variants and percentage of the total amount of genetic variants is indicated.

**Table S4.** Overview of sampling sites at the Veluwe, number of sampled plants per site and coordinates.

| Site | Abbreviation | Sample size | Latitude | Longitude |
| --- | --- | --- | --- | --- |
| Akker Reijerskamp 1 | AR1 | 3 | 52.01481667 | 5.787488889 |
| Akker Reijerskamp 3 | AR3 | 2 | 52.01659167 | 5.789366667 |
| Oud Reemst | OR | 6 | 52.04151667 | 5.8092333 |
| Reijerskamp | REIJ | 3 | 52.016344444 | 5.7723667 |
| Telefoonweg 1 | TW1 | 4 | 52.00221667 | 5.75395 |
| Mossel | MO | 6 | 52.05981667 | 5.75215 |
| Nieuw Reemst | NR | 4 | 52.04274167 | 5.775575 |
| Mossels Veld | MV | 5 | 52.07188333 | 5.736675 |
| Dennenkamp | DK | 6 | 52.02858333 | 5.798766667 |
| Akker Reijerskamp 2 | AR2 | 6 | 52.015675 | 5.788108333 |
| Telefoonweg 2 | TW2 | 6 | 51.998875 | 5.752216667 |

**Figure S1.**
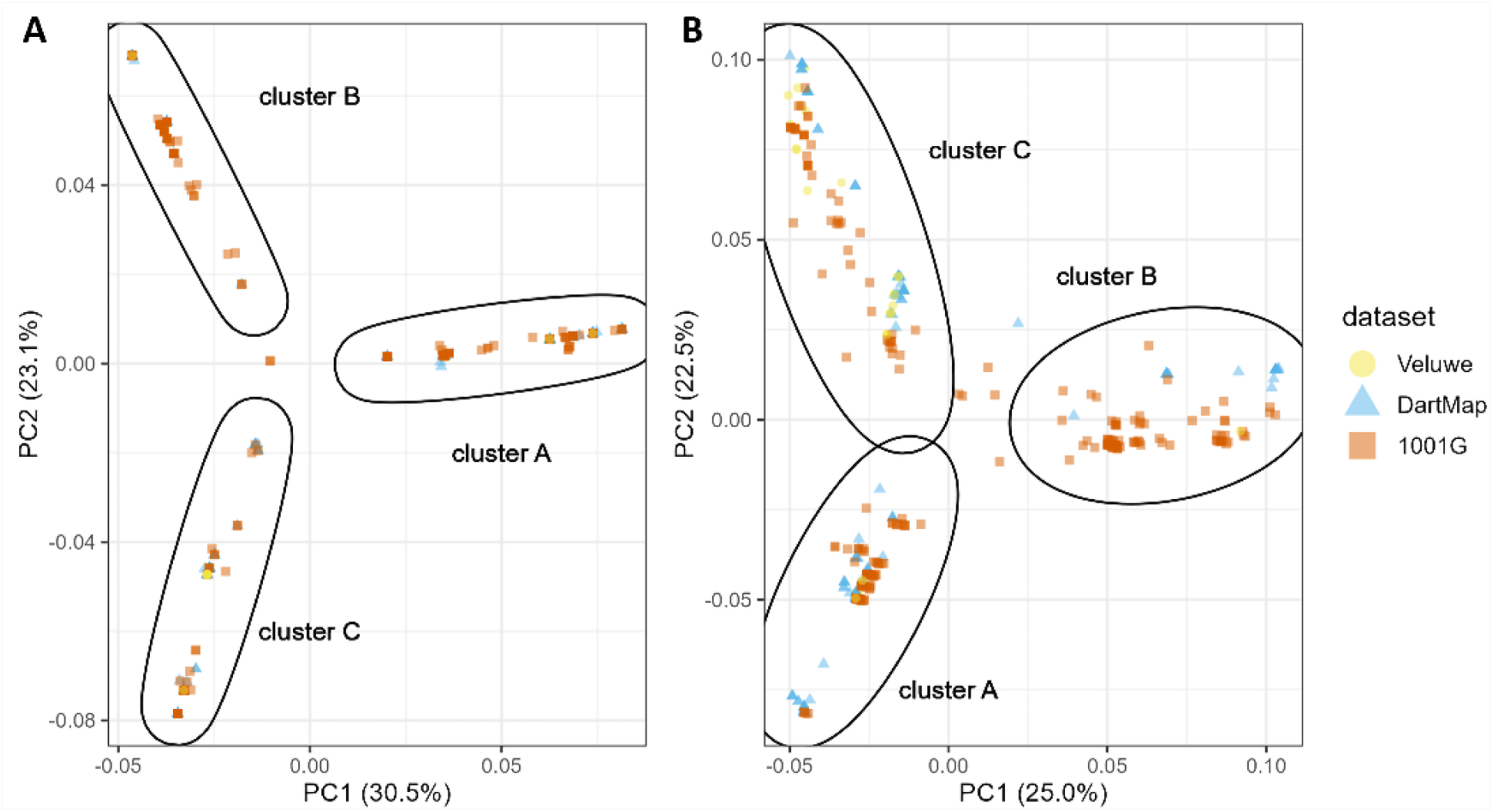
Principal component analysis of organellar genomes. (**A**) Chloroplast genetic variation. (**B**) Mitochondrial genetic variation. Each data point indicates an accession (n = 473). Colours and shapes indicate dataset (Veluwe n = 51, DartMap n = 192, 1001G n = 230). Ellipses indicate organellar clusters based on hierarchical clustering (A, B or C).

**Figure S2.**
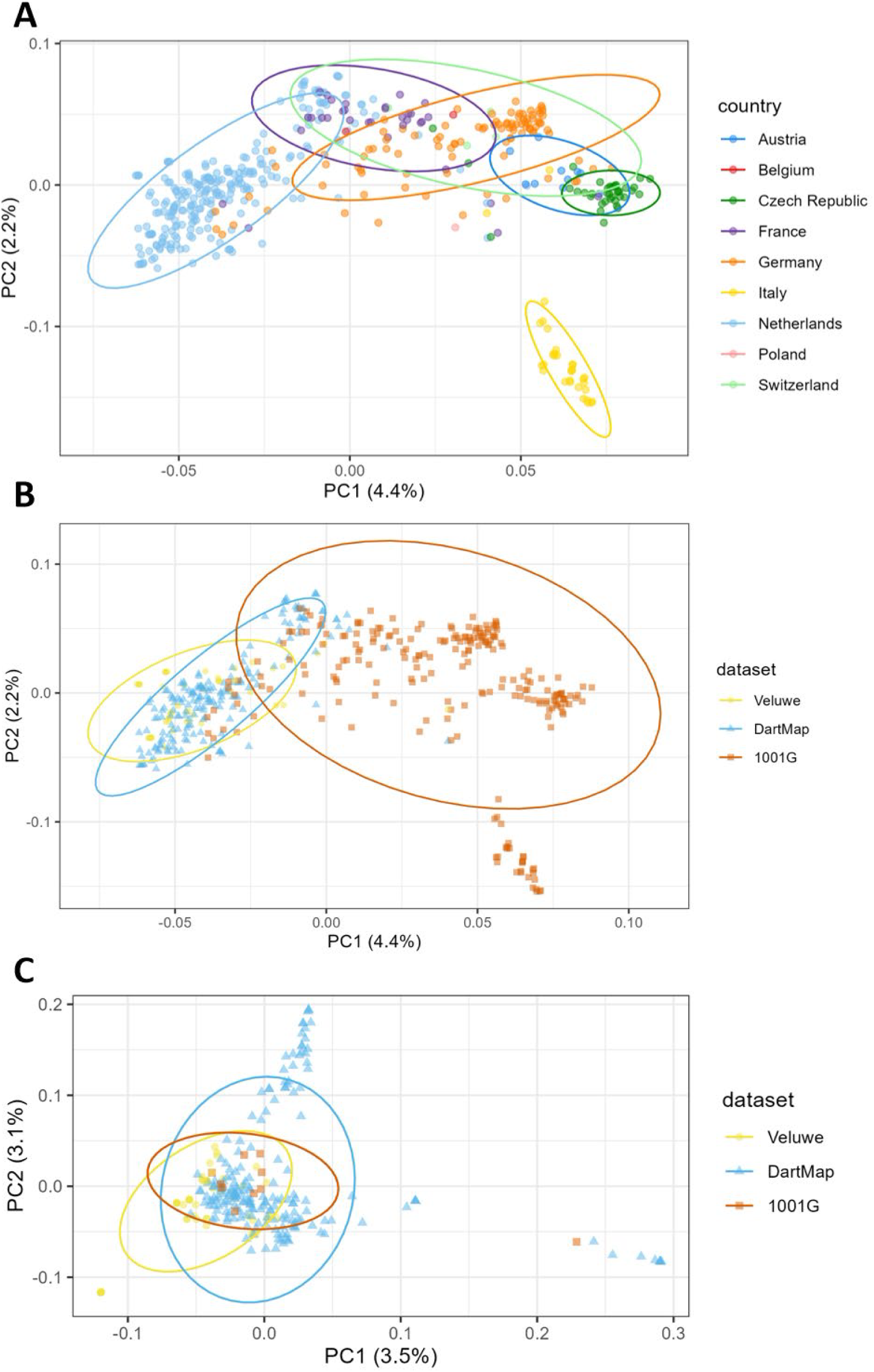
Principal component analysis to assess nuclear population structure. (**A**) Coloured based on country of sampling. Each data point indicates an accession (n = 473). Colours indicate countries and ellipses indicate 95% confidence intervals for countries with more than one accession. (**B**) Coloured based on dataset. Each data point indicates an accession (n = 473). Colours and shapes indicate dataset (Veluwe n = 51, DartMap n = 192, 1001G n = 230) and ellipses indicate 95% confidence intervals. (**C**) Principal component analysis of nuclear genome of accessions originating from the Netherlands. Each data point is an accession (n = 253). Colours and shapes indicate the dataset (Veluwe n = 51, DartMap n = 192, 1001G n = 10). Ellipses indicate 95% confidence intervals.

**Figure S3.**
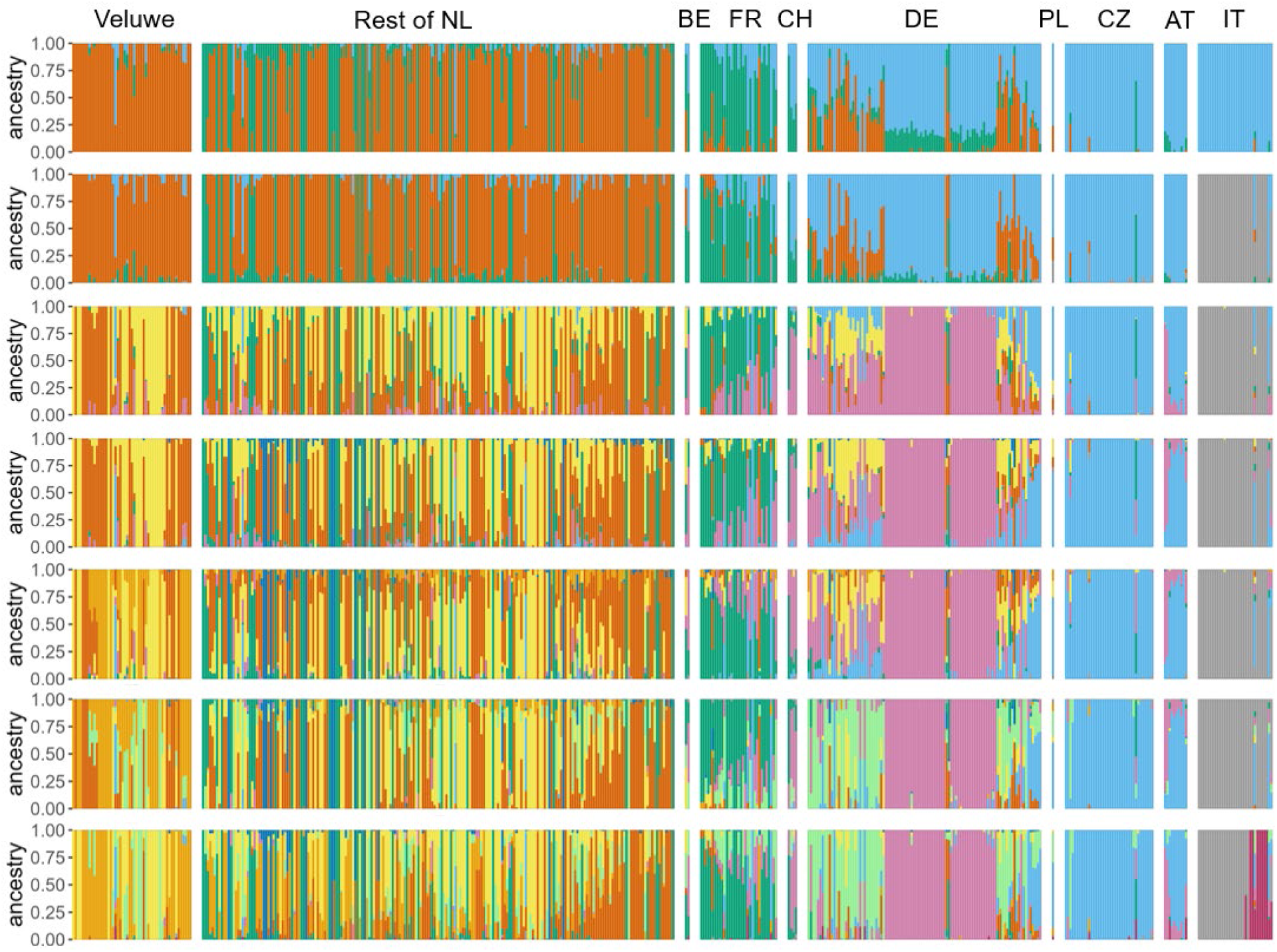
Admixture analysis results for ancestry estimation of the nuclear genome. Y-axis indicates proportion of shared ancestry. K = 3, 4, 6, 7, 8, 9 and 10 (number of ancestral clusters) are displayed from top to bottom. Individual bars are individual accessions grouped according to country. Countries are organized from west (left) to east (right) and within countries accessions are ordered according to longitude (west left and east right). Accessions from the Netherlands are separated between Veluwe dataset (n = 51) and the rest of Dutch accessions (n = 202). BE is Belgium (n = 2); FR is France (n = 33); CH is Switzerland (n = 4); DE is Germany (n = 100); PL is Poland (n = 1); CZ is Czech Republic (n = 38); AT is Austria (n = 10), and IT is Italy (n = 32). Colours indicate ancestral clusters.

**Figure S4.**
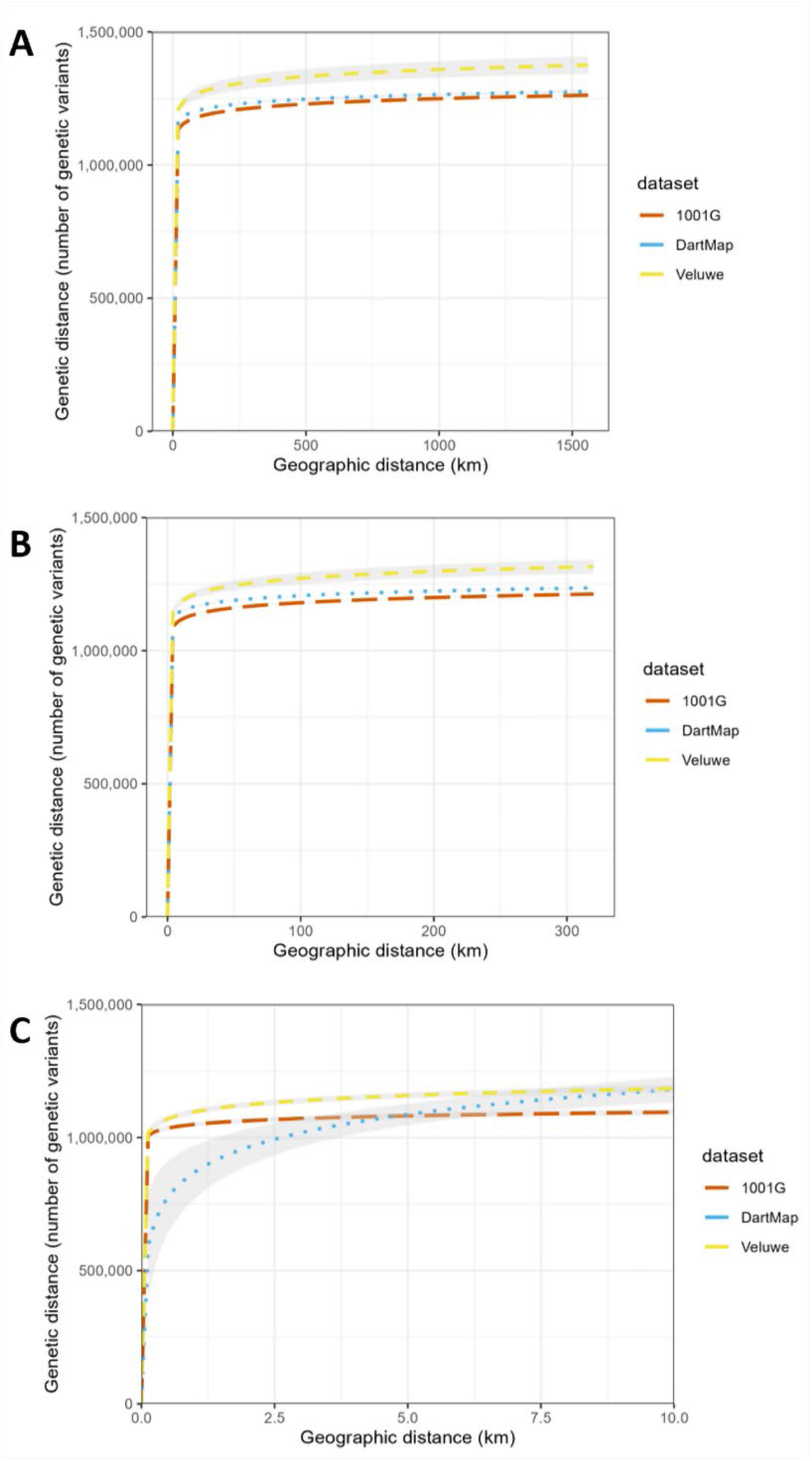
Logarithmic models of genetic versus geographic distances per dataset. (**A**) Plot comparing the logarithmic model of each dataset (1001G, DartMap or Veluwe) on a geographic distance up to 1500 km. (**B**) Plot comparing the logarithmic model of each dataset (1001G, DartMap or Veluwe) on a geographic distance up to 300 km. (**C**) Plot comparing the logarithmic model of each dataset (1001G, DartMap or Veluwe) on a geographic distance up to 10 km. Line colour and type indicates dataset and grey shade indicates standard error. X-axis display pairwise geographic distance between accessions, expressed in kilometre. Y-axis displays pairwise genetic distance between accessions, expressed as number of genetic variants.

**Figure S5.**
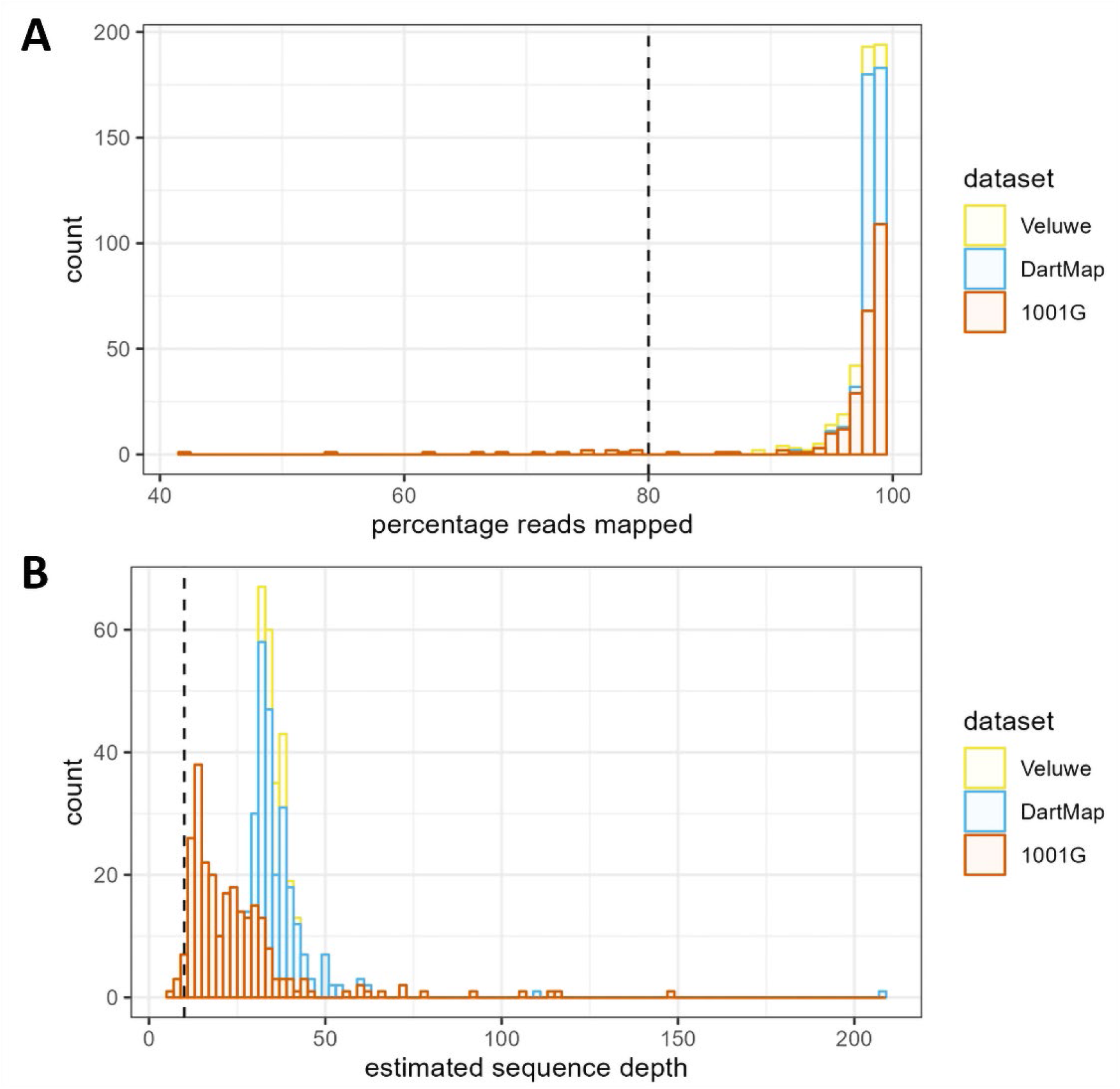
Alignment metrics of nuclear genome per dataset of variant calling pipeline. (**A**) Histogram showing sequencing coverage by percentage reads mapped of the nuclear genome and vertical line indicates cut-off of minimal 80% reads mapped. (**B**) Histogram with estimated reading depth after alignment of the nuclear genome and vertical line indicates cut-off of minimal 10x coverage. Colours indicate datasets. Histogram bars are stacked.

**Figure S6.**
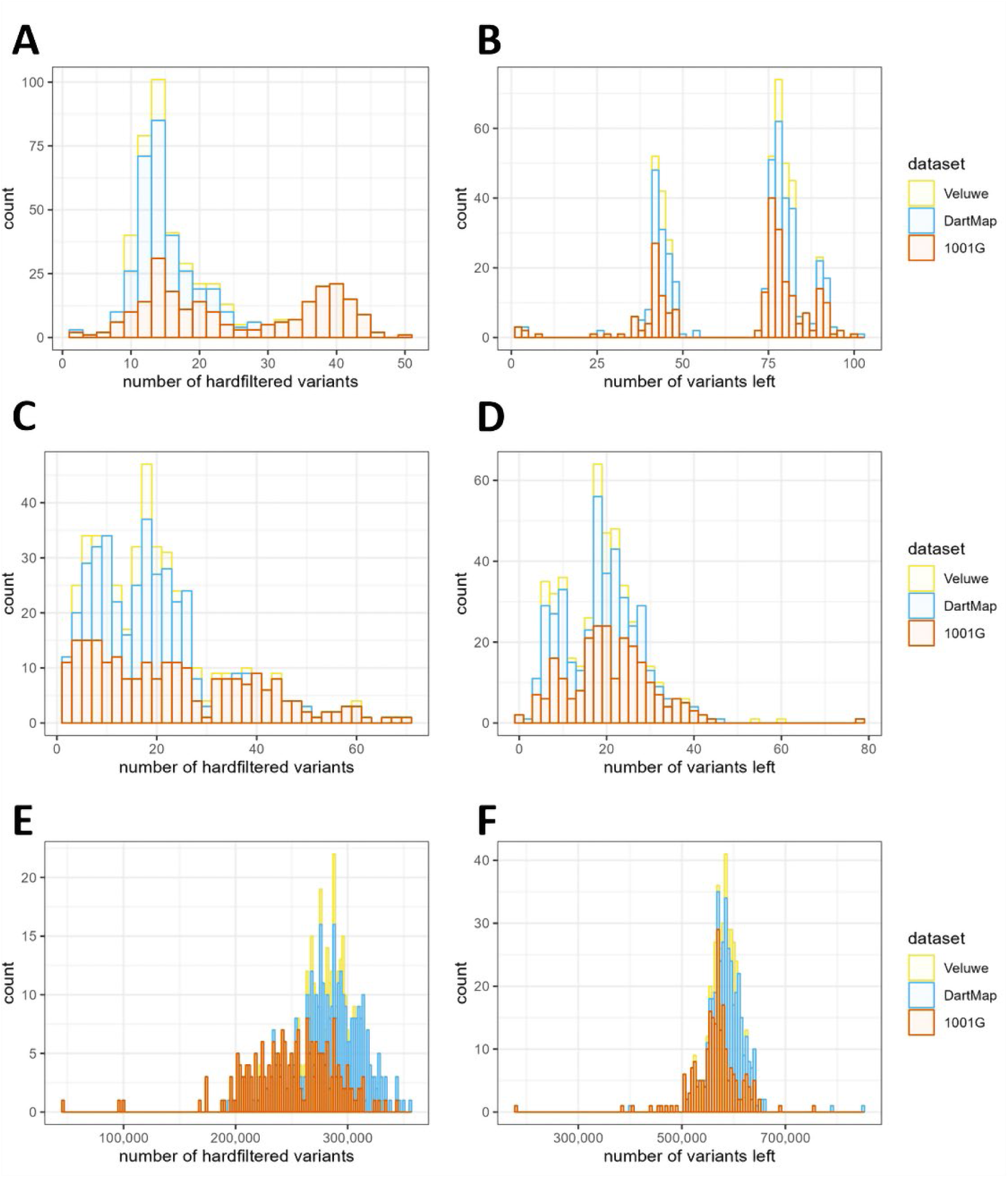
Information on the hard-filtering of chloroplast, mitochondrial and nuclear genetic variants. (**A**) Histogram of number of genetic variants hard-filtered in chloroplast genome. (**B**) Histogram of number of genetic variants left after hard-filtering in chloroplast genome. (**C**) Histogram of number of genetic variants hard-filtered in mitochondrial genome. (**D**) Histogram of number of genetic variants left after hard-filtering in mitochondrial genome. (**E**) Histogram of number of genetic variants hard-filtered in nuclear genome. (**F**) Histogram of number of genetic variants left after hard-filtering in nuclear genome. Colours indicate dataset. Histogram bars are stacked.

